# Anti-thrombotic treatment enhances antibiotic efficiency in a humanized model of meningococcemia

**DOI:** 10.1101/2022.01.10.475613

**Authors:** Jean-Philippe Corre, Dorian Obino, Pierre Nivoit, Aline Yatim, Taliah Schmitt, Guillaume Duménil

**Author notes:** Corresponding authors: G.D. and D.O. The authors equally contributed to this work (Jean-Philippe Corre, Dorian Obino). Institut Curie, PSL Research University, INSERM U932, F-75005 Paris, France.

## Abstract

Meningococcal infections remain particularly difficult to treat. Despite antibiotic therapy, the state of the patients often rapidly deteriorates. Early clinical studies suggest that meningococci acquire a form of resistance to antibiotic treatments during infections. Taking advantage of a humanized animal model of infection, we confirm that adherent bacteria become highly resistant to antibiotic treatments as early as 3-6 hours post infection, although fully sensitive *in vitro*. Within this time frame, meningococci adhere to the endothelium via their type IV pili, proliferate and eventually fill the vessel lumen. Using intravital imaging, we show that rapidly upon infection blood flow is dramatically decreased, thus limiting antibiotic access to infected vessels. Concomitantly, fibrin is deposited inside infected vessels in proximity to bacterial aggregates. Pharmacologically impairing thrombin generation by inhibiting Factor X activity not only improves blood flow in infected vessels, but also enhances the efficacy of the antibiotic treatment. Our results indicate that the combined administration of anticoagulants together with antibiotics might represent a therapeutic approach to treat meningococcal sepsis more efficiently.

## Introduction

Despite intensive medical care meningitis and *purpura fulminans* induced by *Neisseria meningitidis* remain a major concern worldwide, reflecting our limited understanding of the mechanisms underlying these diseases. Systemic forms of the infection are often the most life-threatening as they account for up to 90% of the mortality associated to invasive meningococcal diseases (IMD) (Brandtzaeg & van Deuren, 2012; van de Beek *et al*, 2006). Meningococcal sepsis is characterized by specific clinical manifestations such as the appearance of skin purpuric/petechial lesions and most importantly a particularly fast and severe progression (Stephens, 2007; Thompson *et al*, 2006). Otherwise healthy patients evolve from an absence of symptoms to being critically ill within 12 to 24 hours. Although made difficult by the relatively unspecific clinical features at disease onset, the early recognition and treatment of meningococcal systemic infections by parenteral administration of antibiotics is key in determining patient outcome (Dellinger *et al*, 2013). Remarkably, despite antibiotic administration, disease often progresses relentlessly towards septic shock making these infections particularly difficult to treat. While it is generally considered that treatments will be inefficient past a certain tipping point, the reasons for these failures remain largely unexplained. A possible explanation would be that bacteria acquire a form of resistance to the treatments at a certain point of infection. Accordingly, biopsies of skin lesions performed on patients suffering from meningococcal septic shock revealed the presence of live bacteria up to 13 hours after the initiation of the antibiotic treatment (de Greeff *et al*, 2008; van Deuren *et al*, 2000; van Deuren *et al*, 1993). The mechanisms underlying bacterial antibiotic resistance at play during meningococcal infections remains elusive. Due to the human-species specificity of the bacterium, the lack of reliable animal models of meningococcal infections has precluded any in-depth experimental studies of this process. Today, this limitation has been overcome by the development of a human skin xenograft mouse model that supports the anastomosis of dermal human blood vessels with the murine circulatory system (Melican *et al*, 2013). This model recapitulates the hallmark features of meningococcal systemic infections, such as the adhesion of bacteria to the human endothelial surface and the occurrence of skin purpuric lesions, hence allowing the exploration of *Nm* systemic infections in the full complexity of the host tissue *in vivo* (Bonazzi *et al*, 2018; Melican *et al*., 2013). Combining the use of such a model and intravital imaging, we show that bacteria become resistant to antibiotic treatments as early as 3-6h post infection and that this effect is due to a fast decrease in blood flow induced by the deposition of fibrin inside infected vessels.

## Results

### Meningococci adhering along the vascular wall rapidly acquire antibiotic resistance

We first sought to evaluate whether the humanized model of meningococcal infection could provide new information on the clinical observations pertaining to antibiotic resistance during infection. In this model, intravenously injected *Neisseria meningitidis* circulate in the blood stream and a fraction of bacteria specifically colonizes dermal human vessels in the grafted skin, forming tight bacterial aggregates that progressively fill the infected human vessels (Fig 1A-B) (Manriquez *et al*, 2021; Melican *et al*., 2013). Bacterial sensitivity to intravenously injected antibiotics was assessed in the animal model 3h, 6h and 16h post infection by treating mice with Cefotaxime (CTX, Fig 1C), a standard of care antibiotic for meningococcal infections in patients. Mice were treated with doses reflecting the concentration used in patients (200 mg/kg) (Griffiths *et al*, 2018). As for most clinical meningococcal isolates, the strain used for these experiments is sensitive to low doses of Cefotaxime when cultured in medium containing serum *in vitro* (Fig EV1). Antibiotic administration led to a strong reduction in the numbers of blood-circulating bacteria regardless the time post infection (Fig 1D). Numbers of adherent bacteria also decreased upon antibiotic treatment 3h post infection. In contrast, 6 and 16h post infection, bacteria became strongly resistant to the antibiotic treatment. While bacterial counts were reduced by up to 11 000-fold 3h post infection (55 410 ± 12 445 versus 5 ± 3 CFU/mg of skin in non-treated versus CTX-treated animals, respectively), 16h post infection bacterial counts only decreased by 12-fold (90 731 ± 50 772 versus 7 642 ± 2 200 CFU/mg of skin in non-treated versus CTX-treated animals, respectively) (Fig 1E). Similar results were obtained at 6h post infection (Fig 1E). Taken together, these data show that between 3h and 6h post infection, bacteria adhering to the endothelium experience a change, either intrinsic or in their environment, which leads to antibiotic treatment resistance. Given the ability of meningococci to occlude infected vessels combined with their pro-coagulant activity, a straightforward hypothesis would be that a reduction in microvascular patency could alter the access of antibiotics to bacteria.

**Figure 1.**
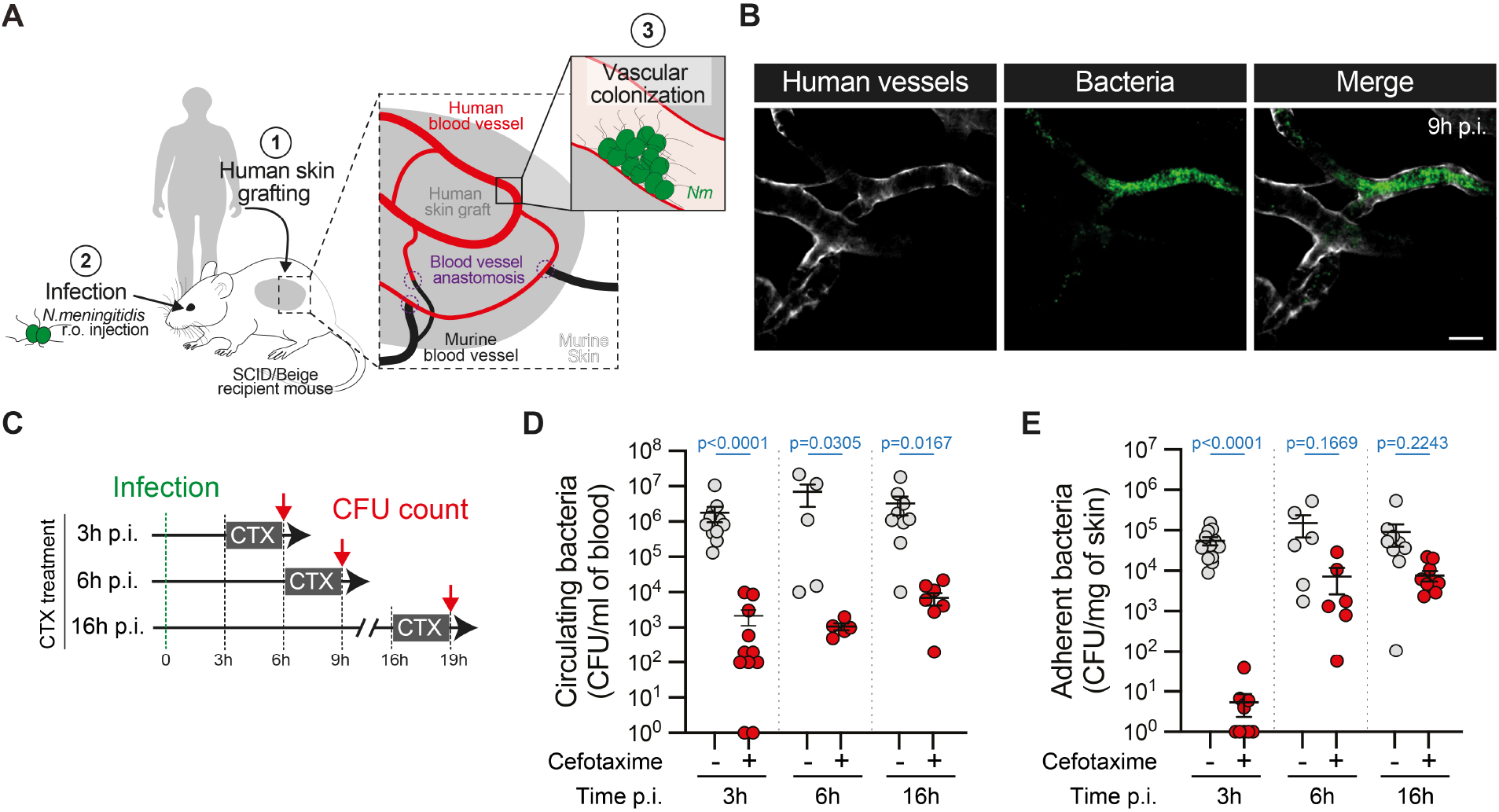
Limited antibiotic efficiency during *Neisseria meningitidis* vascular colonization. **A** Schematic representation of the human skin xenograft mouse model. **B** Representative whole-mount immunofluorescence pictures (maximum intensity z-projection) of the colonization of human vessels (grey, anti-human Collagen IV) by mCherry-expressing *Neisseria meningitidis* (green) in grafted animals 9 hours post infection (p.i.). Scale bar: 40 μm. (n = 3 mice). **C** Schematic representation of the experimental approach used to assess bacterial sensitivity to Cefotaxime (CTX) treatment at 3h, 6h and 16h post infection (p.i.). **D, E** Bacterial colony forming unit (CFU) counts from **(D)** blood (circulating bacteria) and **(E)** dissociated human xenografts (adherent bacteria) collected from mice infected for the indicated time points and treated (+) or not (-) with Cefotaxime (CTX). (n = 12, 6 and 10 mice per group at 3h, 6h and 16h p.i., respectively, pooled from N = 3 independent experiments per time point). Data information: In (D-E), data are presented as the mean ± SEM. Kruskal-Wallis test with Dunn’s correction for multiple comparisons.

### Meningococcal vascular colonization leads to a rapid and complete block of blood flow

Using intravital imaging, we next assessed whether meningococcal vascular colonization alters vascular perfusion and consequently the proper access of antibiotics to sites of infection. Blood flow velocities were assessed using 1 μm fluorescent microspheres injected intravenously. High temporal resolution imaging with a spinning disk confocal microscope allowed the visualization of beads freely circulating in human vessels of non-infected animals (Fig 2A and Movie EV1). Beads circulated at average speeds of 1174 ± 141 μm/s in the observed arterioles (n = 31, average diameter 35 μm) and 505 ± 219 μm/s in venules (n = 13, average diameter 54 μm). Similar values were observed for the first 2 hours of infection (Fig 2B). In contrast, a steep reduction was observed at 3 hours post infection in all infected vessels, culminating in a complete absence of flow at 4 hours post infection (Fig 2B and Movie EV1). No variation in the blood vessel diameter was observed during the infection (Fig 2C), indicating that blood flow reduction was not due to a global collapse of vessels. We recently showed that meningococci can infect dermal arterioles and venules (Manriquez *et al*., 2021) and the same dramatic decrease in blood flow was observed in infected venules and arterioles (Fig 2D), suggesting that the effect observed is independent of the vessel type. Taken together, our results indicate that by preventing the perfusion of infected vessels starting at 3 hours post infection, vascular colonization by meningococci limits the bioavailability of antibiotics, thus restraining their ability to eliminate intravascular adherent bacteria. We then explored whether this blood flow reduction was due to the bacteria themselves or secondary to intravascular coagulation or thrombosis.

**Figure 2.**
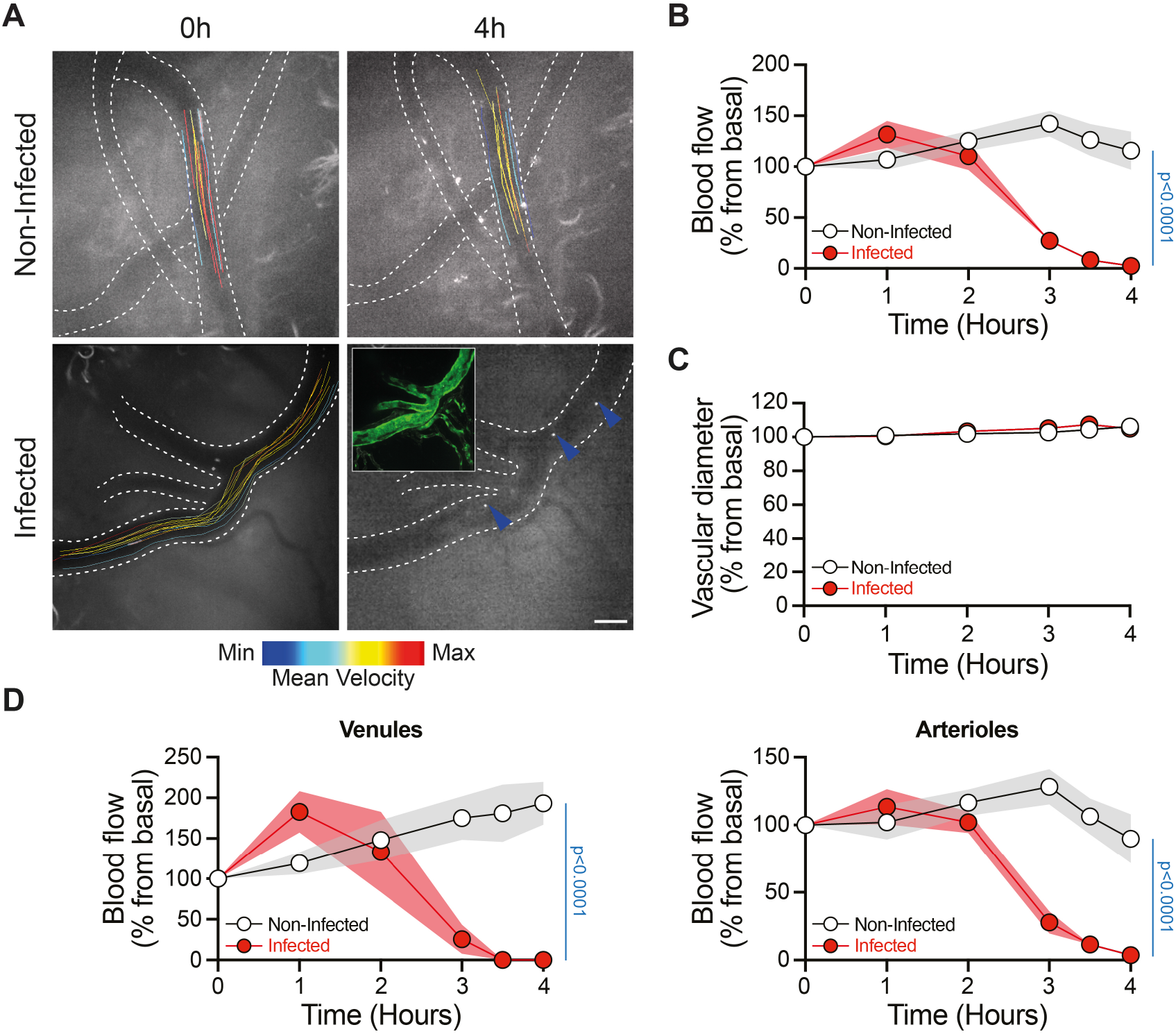
Vascular colonization by *Neisseria meningitidis* blocks the blood flow. **A** Representative image of fluorescent microsphere tracking as a read-out of blood flow during the early phase of the infection (0h and 4h p.i.). Tracks were color-coded according to the mean velocity of the corresponding microsphere. Inset shows the extent of vascular colonization by iRFP-expressing *Nm* (green) 4h post infection. Blue arrowheads indicate immobilized microspheres. Scale bar: 50 μm. **B, C** Normalized blood flow **(B)** and vessel diameter **(C)** as a function of the time post infection. (n = 18 and 19 vessels for non-infected and infected animals, respectively, pooled from N = 2 mice per condition and imaged independently). **D** Data in (B) were classified according to the vessel type: venules (left) and arterioles (right). (n = 13 and 14 arterioles, and 5 and 5 venules in non-infected and infected mice, respectively). Data information: In (B-D), data are presented as the mean ± SEM. Two-way ANOVA test.

### Platelets are not involved in the acquisition of antibiotic resistance

Meningococcal systemic infections are associated with disturbed coagulation ranging from vessel occlusion and thrombus formation to disseminated intravascular coagulation (DIC), as evidenced by the post-mortem analysis of skin biopsies (Harrison *et al*, 2002; Pathan *et al*, 2003; Sotto *et al*, 1976). Similar observations have been recently made using the humanized mouse model of meningococcal infections (Manriquez *et al*., 2021). We therefore tested whether infection-induced microvascular thrombosis might restrict the proper access of antibiotics to adherent meningococci.

We first assessed the role of platelets in the process. Visualization of platelets by intravital imaging showed limited platelet aggregation/accumulation, if any, at sites of bacterial adhesion (Fig EV2A). Platelets could be found in spaces where no bacteria were found, and small volumes of blood expected. Interestingly, no evidence of bacteria-platelet interactions could be found, in contrast to what has been observed with other pathogenic bacteria (Fitzgerald *et al*, 2006; Sun, 2006). We observed circulating platelets moving past bacterial aggregates without evidence of adhesive contacts (Movie EV2). We then evaluated the impact of platelet depletion on the efficiency of Cefotaxime treatment (Fig EV2B). To this end, grafted mice were administered an antibody targeting the platelet receptor GPIbα/CD42b 6h prior to the infection, as previously described (Bergmeier *et al*, 2000; Nieswandt *et al*, 2000). This treatment led to the efficient depletion of platelets (Fig EV2C). Antibiotic treatment was then administrated 16h post infection and bacterial counts assessed 3 hours later. As expected, low numbers of blood-circulating bacteria were found upon antibiotic treatment when compared to the infected but non-treated control grafted animals (Fig EV2D). Numbers of adherent bacteria following the antibiotic treatment were similar to the numbers observed in non-depleted control animals (6 835 ± 3 495 versus 7 642 ± 2 200 CFU/mg of skin in platelet-depleted versus non-depleted animals, respectively) (Fig EV2E to be compared to Fig 1E), suggesting that platelets do not limit the efficacy of antibiotic treatment upon vascular colonization by *Neisseria meningitidis*. Taken together, these results argue against a role for platelets in restricting the efficiency of antibiotic treatments upon meningococcal systemic infections.

### Coagulation inhibition restores antibiotic efficiency

Clinical and experimental evidence point to the presence of fibrin clots in the vicinity of infected vessels (Faust *et al*, 2001; Guarner *et al*, 2004; Melican *et al*., 2013). Using intravital imaging, we therefore monitored the deposition of fibrin inside infected vessels during vascular colonization by *Neisseria meningitidis*. In the first 3 hours following infection, no evidence of fibrin deposition could be found. In contrast, starting 3 hours post infection we observed the progressive accumulation of fibrin close to sites of bacterial adhesion (Fig 3A-B, Fig EV3A and Movie EV3). In absence of infection, fibrin deposition was not observed (n = 17 vessels). Thrombus formation thus coincides with the time of blood flow decrease and the emergence of the resistance to the antibiotic treatment (Fig 1E and Fig 2B). Importantly, fibrin deposition occurs in most infected vessels, reaching 64% coverage at 4h30 post infection (Fig 3B). Interestingly, in patients, circulating monocytes have been shown to express Tissue Factor (TF) during meningococcemia (Osterud & Flaegstad, 1983), suggesting a role for TF in triggering disseminated intravascular coagulation (DIC). Accordingly, TF expression on the surface of monocytes (CD115^+^ Gr-1^+^) from infected mice increased with kinetics similar to microvascular fibrin deposition (Fig EV4A). In contrast, endothelial cells did not show any increase in TF expression, even though E-Selectin expression strongly increased, as shown previously (Manriquez *et al*., 2021) (Fig EV4B). We next sought to determine whether infection-induced fibrin deposition negatively impacts the efficacy of antibiotic therapy by inhibiting the thrombin-dependent cleavage of fibrinogen into fibrin strands. Fondaparinux prevents the formation of fibrin clots by inhibiting Factor Xa, and thus the conversion of prothrombin into active thrombin (Bauer *et al*, 2002). To ensure the constant delivery of the anticoagulant treatment to grafted mice, Fondaparinux was infused subcutaneously using osmotic minipumps. Grafted mice were then infected, and the antibiotic treatment (Cefotaxime, CTX) was administrated 16h post infection (Fig 3C). As expected, Fondaparinux strongly reduced plasma Factor Xa (Fig 3D). Interestingly, Fondaparinux treatment *per se* did not alter, nor promote meningococci survival either in the blood or adhering to the vessel walls (Fig EV3B-C). The antibiotic treatment also efficiently decreased the numbers of circulating bacteria in conditions of Factor X inhibition (Fig 3E).

**Figure 3.**
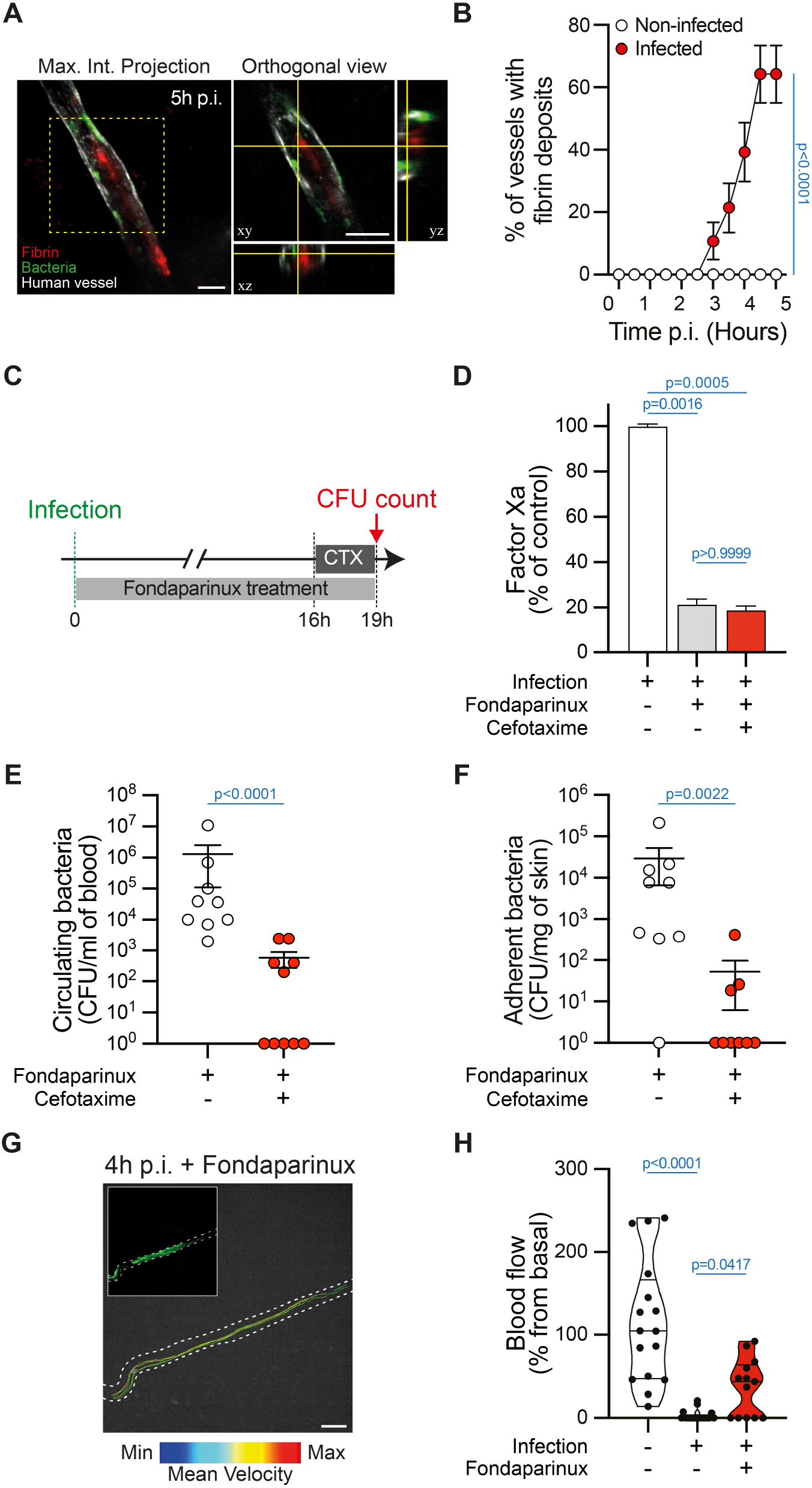
The efficiency of antibiotic treatments is restored upon inhibition of coagulation. **A** Representative image (maximum intensity z-projection) and orthogonal view of fibrin deposition (red, anti-Fibrin) in human vessels (grey, UEA1 lectin) infected by iRFP-expressing *Neisseria meningitidis* (green). Scale bar: 50 μm. (n = 28 infected vessels, pooled from N = 3 mice imaged independently). **B** Percentage of vessels with fibrin deposition in control and infected animals (n = 28 infected vessels and 17 non-infected control vessels, pooled from N = 3 and 2 mice, respectively, imaged independently). **C** Schematic representation of the experimental approach used to assess the impact of coagulation inhibition on bacterial sensitivity to Cefotaxime (CTX) treatment. **D** Chromogenic-based quantification of the coagulation Factor Xa in the plasma of control (no Fondaparinux, no Cefotaxime), Fondaparinux-treated (no Cefotaxime) and Fondaparinux+CTX-treated mice. (n = 8, 10 and 10 mice for control, Fondaparinux-treated, and Fondaparinux+CTX-treated mice, respectively, pooled from N = 3 independent experiments). **E, F** Bacterial colony forming unit (CFU) counts from **(E)** blood (circulating bacteria) and **(F)** dissociated human xenografts (adherent bacteria) collected from mice treated or not with the Fondaparinux anticoagulant and infected for 19h and treated or not with Cefotaxime (CTX). (n = 9 mice per group, pooled from N = 3 independent experiments). **G** Representative images (maximum intensity z-projection) of fluorescent microsphere tracking in human vessels (white dashed line) infected by iRFP-expressing bacteria (green, inset) in mice treated with Fondaparinux. Tracks were color-coded according to the mean velocity of the corresponding microsphere. Scale bar: 50 μm. (n = 3 mice). **H** Violin plot showing the distribution of the blood flow in vessels from non-infected, infected, and infected+Fondaparinux-treated animal 4h post infection. (n = 16, 19 and 13 vessels for non-infected, infected, and infected+Fondaparinux, respectively, pooled from N = 2, 2 and 3 mice for control, infected, and infected+Fondaparinux-treated animals, respectively). Data information: In (D), data are presented as the mean ± SEM. Kruskal-Wallis test with Dunn’s correction for multiple comparisons. In (E-F), data are presented as the mean ± SEM. Two-tailed Mann-Whitney test. In (H), data are presented as violin plot showing the distribution of the dataset. Kruskal-Wallis test with Dunn’s correction for multiple comparisons.

Strikingly, upon inhibition of coagulation and antibiotic treatment, numbers of adherent bacteria in the human grafts were decreased by more than 500-fold when compared to mice that did not received antibiotics (29 591 ± 23 065 versus 52 ± 45 CFU/mg of skin in Fondaparinux-versus Fondaparinux+CTX-treated animals, respectively) (Fig 3F). Accordingly, the perfusion of infected vessels was partially restored upon inhibition of coagulation by Fondaparinux (Fig 3G-H and Movie EV4). Taken together these results strongly suggest that meningococcal-induced fibrin deposition reduces microvascular patency and thereby the tissue distribution of antibiotics, thus preventing their efficacy.

## Discussion

Our results show that within a few hours of infection, meningococcal vascular colonization leads to a dramatic reduction in the perfusion of infected vessels. While bacterial colonization alone is sufficient to partially occlude vessels and reduce blood flow, ensuing thrombus formation leads to the complete cessation of blood flow in affected vessels. This, in turn, has important implications for disease progression. Complete absence of perfusion will result in severe ischemia and irreversible tissue injury, driving the rapid progression of the disease towards multiorgan failure. Another consequence of reduced microvascular perfusion, detailed in this study, is its impact on the access of antibiotics to target tissues. This is observed in the animal model described here but also in human cases where bacteria have been demonstrated to resist intensive antibiotic treatment (van Deuren *et al*., 1993).

The molecular process driving thrombin generation during meningococcal infection has been extensively studied clinically and *in vitro* but remains to be fully established (Lecuyer *et al*, 2017). Our results using a humanized model of *Neisseria meningitidis* systemic infection show extensive fibrin deposition in close proximity to bacterial aggregates early during infection and provide new insight into meningococci-induced thrombus formation. Our results suggest that the process is largely independent of platelets, which do not stably interact with meningococci, as opposed to many other bacterial infections (Fitzgerald *et al*., 2006; Sun, 2006). Rather, thrombus formation appears to be triggered by the extrinsic pathway of blood coagulation through the rapid expression of TF/CD142 by monocytes. Accordingly, in meningococcemia cases (Osterud & Flaegstad, 1983) and the humanized model used here, monocytes express TF at early stages of infection. In contrast and despite massive local bacterial accumulation, human endothelial cells did not display TF expression *in vivo*. Interestingly, the contact (intrinsic) pathway of blood coagulation was also shown to be activated during meningococcal infections (Wuillemin *et al*, 1995). The negative charges present on the bacterial LOS or capsule could be a trigger of the contact pathway (Oehmcke & Herwald, 2010). This could provide an alternative explanation for fibrin deposition in proximity to bacterial colonies. In any case, vascular colonization by *Neisseria meningitidis* leads to the rapid formation of thrombi that restrict the efficacy of antibiotic treatment. Many pathogenic bacteria have evolved specific strategies to inhibit coagulation, a process that is thought to favor bacterial dissemination (Popov *et al*, 1991). Meningococci do not seem to be equipped to efficiently block thrombus formation, hence permitting the thrombotic occlusion of infected vessels and the sealing of the infected regions by the host. This likely reflects the main lifestyle of *Neisseria meningitidis* at the surface of the nasopharyngeal epithelium rather than in the vasculature.

From the clinical point of view, our observations suggest that anticoagulation therapy could potentially provide a synergistic effect in conjunction with antibiotic treatment for meningococcal infections. A number of clinical studies have tested the benefits of anticoagulants such as activated protein C in combination with antibiotics. Activated protein C acts as a negative regulator of coagulation by proteolytically inactivating coagulation cofactors Factor Va and Factor VIIIa. Unfortunately, the use of recombinant Protein C has had variable success (Silva *et al*, 2010). Our study suggests that anticoagulation would mainly be effective in the very early stages of infection and that direct targeting of Factor Xa or thrombin may be more effective than Protein C. The efficacy of anticoagulation described here is likely to be linked to the pathogenesis of meningococcal infections and more specifically to the ability of this bacterium to adhere to the endothelium and colonize vessels. The utility of anticoagulation is thus likely to depend on the pathogen. Although the process described here is local, in close association with infected vessels, it is a host-driven response and a likely precursor of DIC. The induction of local coagulation appears to represent a tipping point in the progression of the disease after which antibiotic therapy is no longer as efficient.

While our study suggests the potential of anticoagulant therapy for increased efficacy of antibiotic treatment, it also argues that such therapy would have to be administered early in the progression of the disease and that thrombolytic therapy is likely to be more efficacious once the bacteria have taken hold and thrombi are established. Importantly, the use of anticoagulant and fibrinolytic therapy is always associated with a potential risk of bleeding that will need to be carefully assessed. Although anticoagulation has been used in the context of sepsis in the past (Levi *et al*, 2001; Silva *et al*., 2010), the strong implication of the cerebral vasculature render safety considerations particularly important for meningococcal infections. That said, experience from ischemic stroke shows that such risks can be managed by the use of strict inclusion criteria based on timing and individual risk/benefit assessment.

## Materials and Methods

### Mice

SCID/Beige (CB17.Cg-*Prkdc*^*scid*^*Lyst*^*bg-J*^/Crl) mice were purchased from Charles Rivers (France) and housed under specific pathogen-free condition at Institut Pasteur. Mice were kept under standard conditions (light 07.00-19.00h; temperature 22±1°C; humidity 50±10%) and received sterilized rodent feed and water *ad libitum*. All experiments were performed in agreement with guidelines established by the French and European regulations for the care and use of laboratory animals and approved by the Institut Pasteur committee on Animal Welfare (CETEA) under the protocol code DAP 180022. For all experiments, male and female mice between 9 and 16-weeks of age were used. Littermates were randomly assigned to experimental groups.

### Human skin

Normal human skin was obtained from adult patients (20-60 years old), both males and females, undergoing plastic surgery in the service *de chirurgie reconstructrice et plastique* of Groupe Hospitalier Saint Joseph (Paris, France). In accordance with the French legislation, patients were informed and did not refuse to participate in the study. All procedures were approved by the local ethical committee *Comité d’Evaluation Ethique de l’INSERM* IRB 00003888 FWA 00005881, Paris, France Opinion: 11-048.

### Xenograft model of infection

6-13 weeks old mice, both males and females, were grafted with human skin as previously described (Melican *et al*., 2013). Briefly, mice were anesthetized with a mixture of ketamine hydrochloride (100 mg/kg, Boehringer Ingelheim) and xylazine hydrochloride (8.5 mg/kg, Bayer) and a graft bed of approximately 1–2 cm^2^ was prepared on their flank by removing the mouse epithelium and the upper dermis layer. A human skin graft (200 μm thick) comprising the human epidermis and the papillary dermis was immediately placed over the graft bed. Grafts were fixed in place with surgical glue (Vetbond, 3M, USA) and dressings were applied for 2 weeks. Grafted mice were used for experimentation 3-6 weeks post-surgery when the human dermal microvasculature is anastomosed to the mouse circulation without evidence of local inflammation, as previously described (Melican *et al*., 2013). All efforts were made to minimize suffering.

### *Neisseria meningitidis* strains and mouse infection

Experiments were performed using *N. meningitidis* 8013 serogroup C strain (http://www.genoscope.cns.fr/agc/nemesys) (Rusniok *et al*, 2009). Strains were streaked from -80°C freezer stock onto GCB agar plates and grown overnight in a moist atmosphere containing 5% CO_2_ at 37°C. For all experiments, bacteria were transferred to liquid cultures in pre-warmed RPMI-1640 medium (Gibco) supplemented with 10% FBS at adjusted OD_600nm_ = 0.05 and incubated with gentle agitation for 2 hours at 37°C in the presence of 5% CO_2_. Bacteria were washed twice in PBS and resuspended to 10^8^ CFU/ml in 1x PBS. Prior to infection, mice were injected intraperitoneally with 8 mg of human transferrin (Sigma Aldrich) to promote bacterial growth *in vivo* as previously described (Melican *et al*., 2013). Mice were infected by retro-orbital injection of 100 μl of the bacterial inoculum (10^7^ CFU total). When indicated, mice received a retro-orbital injection of 4 mg Cefotaxime (CTX, #C7912, Sigma-Aldrich) at 3h, 6h or 16h post infection and 3h prior to mouse sacrifice. No influence of mice sex on bacterial colonization have been observed.

### Wholemount immunofluorescent staining

Following mouse infection with mCherry-expressing *Neisseria meningitidis* for 9 hours, the human grafts were harvested and fixed in 4% PFA overnight at 4°C with gentle agitation. Skin samples were then washed and permeabilized/blocked overnight at 4°C with gentle agitation using a solution of 0.3% Triton-X100, 1% BSA, 1% normal goat serum in 1X PBS. The basal lamina of human blood vessels was stained using an AlexaFluor647-conjugated anti-human Collagen IV antibody (#51-9871-82, eBioscience, 1/100) diluted in permeabilization/blocking solution and incubated 3 days at 4°C with gentle agitation. Samples were then extensively washed and tissue transparization and mounting were performed using RapidClear 1.47 (SunJin Lab) according to manufacturer’s recommendations.

### *Neisseria meningitidis* growth curves

Bacterial liquid culture was diluted to an OD_600nm_ of 0.05 and 150 μl of the bacterial suspension was transferred into wells of a 96-well plate. Bacteria were then treated or not with the indicated final concentration of Cefotaxime, each performed in quintuplicates. Plates were placed in a Cytation 5 multimode reader (Biotek) at 37°C with 5% CO_2_ and continuous agitation and optical densities at 600 nm were recorded every 30 minutes for 20 hours.

### Colony-forming units (CFU) enumeration

To assess bacteraemia (blood-circulating bacteria) in infected animals, 10 μl of blood was sampled at the time of sacrifice. Serial dilutions of blood were plated on GCB agar plates and incubated overnight at 37°C and in a moist atmosphere containing 5% CO_2_. Bacterial counts were expressed in colony-forming units (CFU) per ml of blood. To assess the extent of vascular colonization by meningococci (adherent bacteria) following mouse sacrifice at indicated times post infection, tissue biopsies were collected using a sterile dermatological biopsy puncher (approximately 4 mm^2^), weighted and placed in 500 μl 1x PBS. Skin biopsies were dissociated and homogenized using a BeadMill4 homogenizer (Fisher Brand) and serial dilutions of skin homogenates were plated on GCB plates incubated overnight at 37°C and in a moist atmosphere containing 5% CO_2_. Bacterial counts were expressed in colony-forming units (CFU) per mg of skin.

### Platelet depletion

Platelets were depleted in grafted mice by retro-orbital injection of 40 μg of anti-mouse GPIbα (CD42b) (#R300, Emfret Analytics) monoclonal antibodies diluted in 100 μl 1x PBS 6 hours prior to infection, as previously described (Bergmeier *et al*., 2000; Nieswandt *et al*., 2000). The efficiency of platelet depletion was assessed by counting platelets on citrated blood using an automated hemocytometer (SCIL Vet ABC plus) according to manufacturer’s instructions.

### Spinning disk confocal intravital imaging

Intravital imaging of the human xenograft was adapted from (Ho *et al*, 2000) and previously described (Manriquez *et al*., 2021). Briefly, 30 min prior to surgery, mice were injected subcutaneously with buprenorphine (0.05 mg/kg, CEVA) and anesthetized by spontaneous inhalation of isoflurane in 100% oxygen (induction: 4%; maintenance: 1.5% at 0.3 L/min). A flap of skin supporting the human graft was immobilized onto a custom-made heated deck (36°C) and continuously moistened with warmed 1x PBS (36°C). Mouse body temperature was maintained at 37°C using a heating pad and oxygen saturation and heart rate were monitored using the pulse oximetry Physiosuit apparatus (Kent Scientific). Image acquisition was performed using a Leica DM6 FS upright microscope equipped with a motorized stage, a Fluotar 25x/0.95 objective (Leica), and coupled to a Yokogawa CSU-W1 confocal head modified with Borealis technology (Andor). Four laser excitation wavelengths (488, 561, 642, and 730 nm) were used in fast succession and visualized with the appropriate long-pass filters. Fluorescence signals were detected using a sCMOS 2048×2048 pixel camera (Orca Flash v2+, Hamamatsu). Metamorph acquisition software (Molecular devices) was used to drive the confocal microscope.

The human vessels were labelled using Dylight755-conjugated UEA-1 lectin (100 μg, Vector Laboratories), platelets using DyLight488-anti-mouse GPIbβ antibody (0.1 μg, #X488, Emfret Analytics) and fibrin deposits using DyLight488-anti-human/mouse fibrin antibody (8 μg, clone 59D8, Sigma-Aldrich). All antibodies and dyes were injected intravenously 15 min prior to intravital imaging. Five to ten fields of view of interest were selected per animal and time-lapse z-stack series (2-2.5 μm z-step, 50-80 μm range) were captured every 30-60 min for 4-5 hours following the intravenous injection of iRFP-expressing bacteria (10^7^ CFU/100 μl 1x PBS). In mice treated with anticoagulant, Fondaparinux was injected subcutaneously 5 min prior to imaging (80 μl, 1 mg).

The blood flow during vascular colonization was monitored by perfusing 1 μm diameter fluorescent microspheres (Yellow/Green Fluoresbrite carboxylate, Polysciences, 10^7^ microspheres/ml 1x PBS) at a rate of 15 μl/min before each z-stack series and images acquired on a single plane at high speed (50 frames per second, 300 frames). Blood flow was calculated from the vessel diameter and the centerline microsphere mean velocity, according to (Hidalgo *et al*, 2009), and normalized with respect to the basal blood flow. Platelet dynamics during vascular colonization was assessed by fast (50 frames/sec) simultaneous dual-color imaging of platelets and bacteria.

### Anticoagulant treatment

The inhibition of coagulation was achieved by treating grafted mice with the factor Xa inhibitor Fondaparinux (ARIXTRA®, ASPEN) prior to and/or during the course of the infection. For short-term infection (6h and 9h), grafted mice received a subcutaneous injection of 80 μl Fondaparinux (corresponding to 1 mg) immediately after the intravenous injection of bacteria and 3h and 6h post infection. For long-term infection (19h), grafted mice received a subcutaneous injection of 80 μl Fondaparinux (corresponding to 1 mg) immediately after the infection and the constant delivery (infusion) of Fondaparinux was then ensured by ALZET® osmotic pumps (200 μl, 8 μl/h during 24h, #2001D) filled with 200 μl Fondaparinux and placed subcutaneously, according to manufacturer’s instructions. The efficiency of the anticoagulant treatment was assessed using a chromogenic enzymatic method (Biophen Heparin 6 kit, #221006, HYPHEN Biomed), according to manufacturer’s instructions.

### Flow cytometry

To monitor the surface levels of Tissue Factor and E-Selectin on human endothelial cells, biopsies of human xenograft harvested at the indicated time post infection were digested with 0.4 mg/ml Liberase TL (Sigma Aldrich) in CO_2_-independent medium for 60 min at 37°C with gentle agitation (100 rpm). The resulting single-cell suspension was passed through a 70-μm cell strainer (BD Bioscience), washed once with FACS buffer (1× PBS supplemented with 0.5% BSA and 2 mM EDTA) and stained in FACS buffer according to standard protocols. Briefly, human and murine Fc receptors were blocked using Human TruStain FcX (Biolegend, #422302, 1/20) and anti-mouse CD16/CD32 (FcBlock clone 2.4G2, BD Biosciences, #553141, 1/200), respectively, and cells were stained for 30 min at 4°C with the following anti-human antibodies: PerCP-Vio700-conjugated REAfinity recombinant anti-CD31 (Miltenyi Biotec, clone REA730, #130-110-673, 1/20), BV421-conjugated anti-Tissue Factor/CD142 (BD Biosciences, clone HTF-1, #744003, 1/20), and PE-conjugated anti-CD62E/E-Selectin (eBioscience, clone P2H3, #12-0627-42, 1/20).

To monitor the surface levels of Tissue Factor on circulating murine monocytes, blood was harvested at the indicated time post infection and treated with 1x RBC lysis solution (BioLegend). Murine Fc receptors were blocked using anti-mouse CD16/CD32 (FcBlock clone 2.4G2, BD Biosciences, #553141, 1/200) and cells were stained in FACS buffer for 30 min at 4°C with the following anti-mouse antibodies: PacificBlue-conjugated anti-CD45 (Biolegend, clone 30-F11, #103126, 1/100), PE-Cy7-conjugated anti-CD11b (BD Biosciences, clone M1/70, #552850, 1/200), APC-conjugated anti-CD115 (Invitrogen, clone AFS98, #17-1152-82, 1/100), BV650-conjugated anti-Gr-1 (Biolegend, clone RB6-8C5, #108441, 1/50), and PE-conjugated anti-Tissue Factor/CD142 (R&D Systems, polyclonal, #FAB3178P, 1/50).

After staining, cells were washed twice in 1× PBS and fixed for 15 min at 4 °C with 1% paraformaldehyde in 1× PBS. Data were acquired using a CytoFLEX S flow cytometer controlled with the CytExpert software (Beckman Coulter). Data analysis was carried out using FlowJo software v10 (Tree Star).

### Statistics

All graphs and statistical analyses were performed with GraphPad Prism 9 (GraphPad Software). No statistical method was used to predetermine sample size. Kolmogorov–Smirnov test was used to assess the normality of all data sets. Scatter dot plots, bar graphs and values are provided as the mean ± sem. P-values were considered as statistically significant when inferior at 0.05. Statistical details of experiments (sample size, replicate number, statistical significance) can be found in the figures, figure legends and source data file.

## Data availability

This study includes no data deposited in external repositories.

## Acknowledgements

This work was supported by the Integrative Biology of Emerging Infectious Diseases (IBEID) laboratory of excellence (ANR-10-LABX-62), and the VIP European Research Council-starting grant (310790-VIP to G.D.). D.O. was supported by a grant from the Agence Nationale de la Recherche (ANR-18-CE15-0006-MeningoChip to G.D.).

## Author contributions

G.D. designed research and supervised overall data analysis; J.-P.C., P.N., A.Y. and D.O. performed and analyzed experiments; T.S. provided human skin samples; D.O. assembled figures; D.O. and G.D. wrote the manuscript.

## Conflict of interest

The authors declare no conflict interests.

**Figure EV1.**
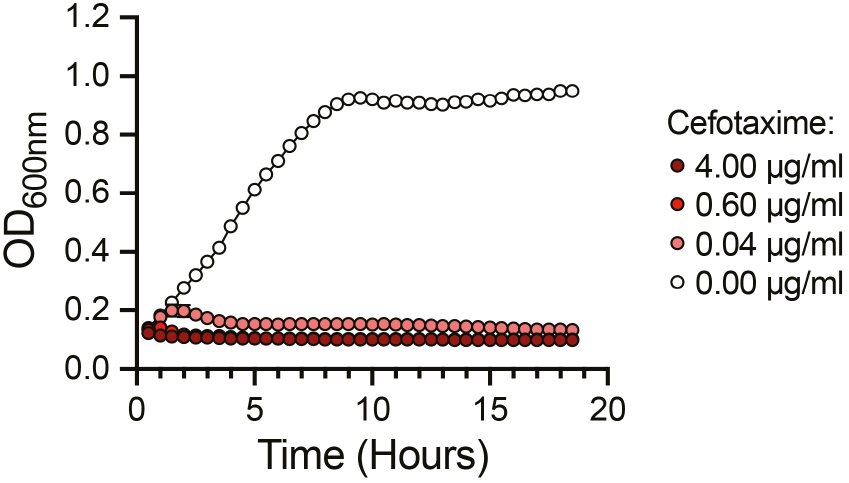
*Neisseria meningitidis* is sensitive to low doses of Cefotaxime *in vitro*. Growth curves of *N. meningitidis* in the presence of the indicated final concentrations of Cefotaxime. (N = 1 experiment). Data are presented as the mean ± SEM of quintuplicates.

**Figure EV2.**
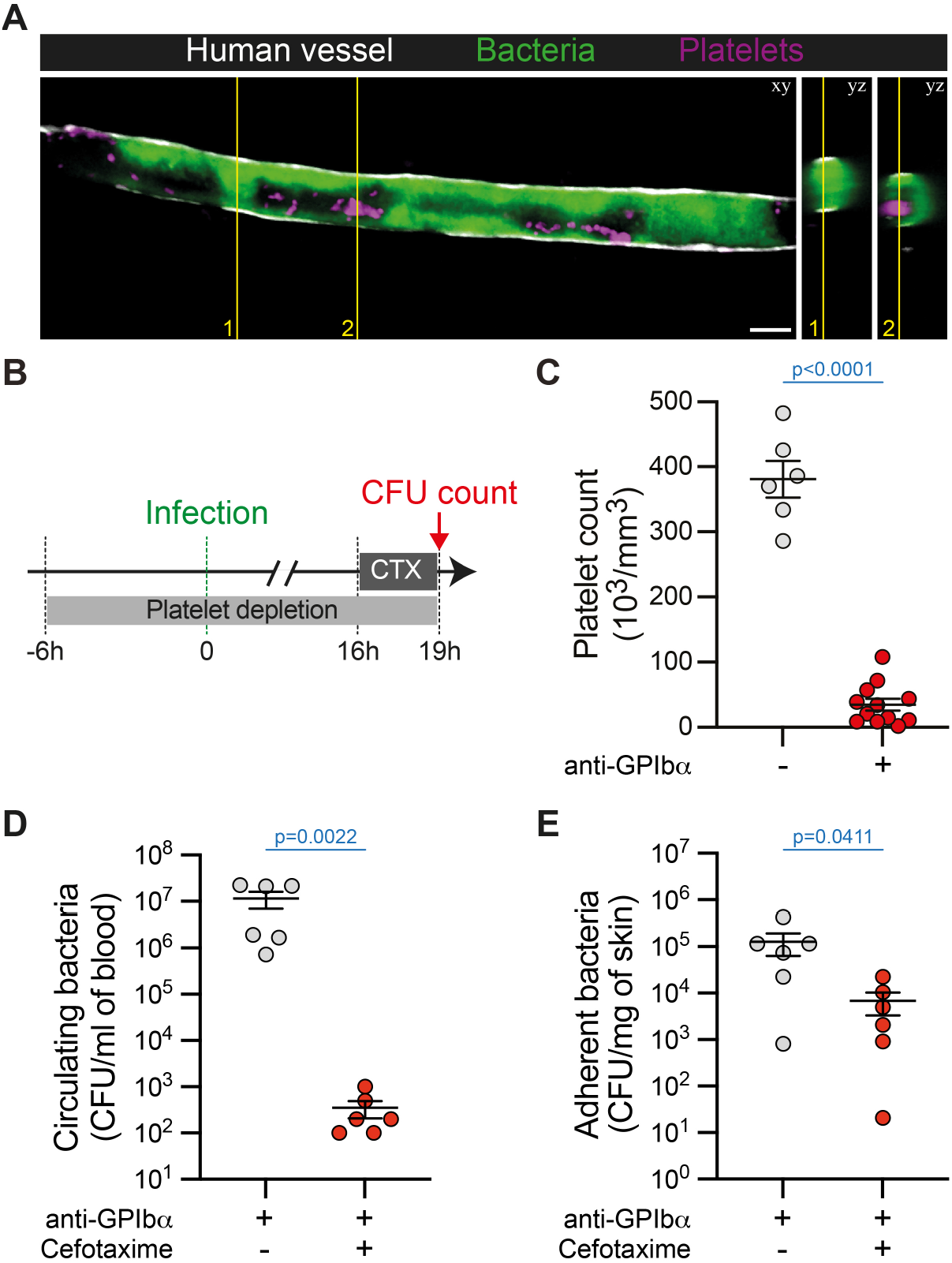
Platelets do not prevent the efficiency of the antibiotic treatment. **A** Representative image (maximum intensity z-projection) and orthogonal view of platelets (purple, anti-GPIbβ) in bacteria-free spaces in human vessels (grey, UEA1 lectin) infected by iRFP-expressing *Neisseria meningitidis* (green) for 4 hours. Scale bar: 25 μm. (n = 3 mice). **B** Schematic representation of the experimental approach used to assess the impact of platelet depletion on bacterial sensitivity to Cefotaxime (CTX) treatment. **C** Quantification of platelet numbers in the blood of mice treated (+) or not (-) with the platelet-depleting antibody directed against GPIbα. (n = 6 and 12 mice for control and anti-GPIbα-treated mice, respectively, pooled from N = 2 independent experiments). **D, E** Bacterial colony forming unit (CFU) counts from **(D)** blood (circulating bacteria) and **(E)** dissociated human xenografts (adherent bacteria) collected from platelet-depleted mice (anti-GPIbα) infected for 19h and treated (+) or not (-) with Cefotaxime (CTX). (n = 6 mice per group, pooled from N = 2 independent experiments). Data information: In (C), data are presented as the mean ± SEM. Unpaired t test. In (D-E), data are presented as the mean ± SEM. Two-tailed Mann-Whitney test.

**Figure EV3.**
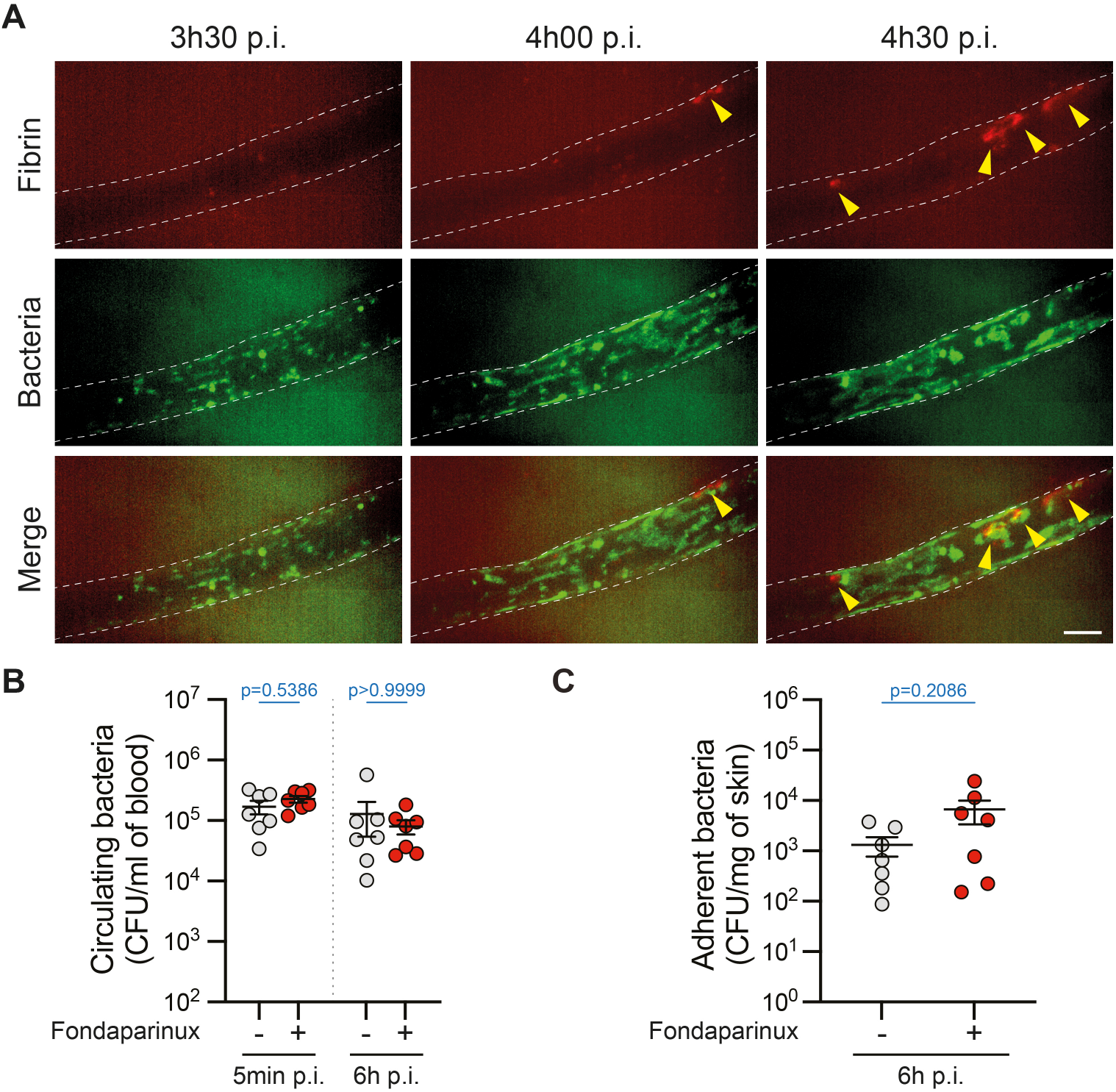
The efficiency of the antibiotic treatment is restored upon coagulation inhibition. **A** Representative time sequence (maximum intensity z-projection) of fibrin deposition (red, anti-Fibrin) in human vessels (dashed line) infected by iRFP-expressing *Neisseria meningitidis* (green). Scale bar: 50 μm. (n = 28 infected vessels, pooled from N = 3 mice imaged independently). **B, C** Bacterial colony forming unit (CFU) counts from **(B)** blood (circulating bacteria) and **(C)** dissociated human xenografts (adherent bacteria) collected from control (-) and Fondaparinux-treated (+) mice infected for 5 minutes or 6 hours. (n = 7 mice per group, pooled from N = 2 independent experiments). Data information: In (B-C), data are presented as the mean ± SEM. (B) Kruskal-Wallis test with Dunn’s correction for multiple comparisons. (C) Two-tailed Mann-Whitney test.

**Figure EV4.**
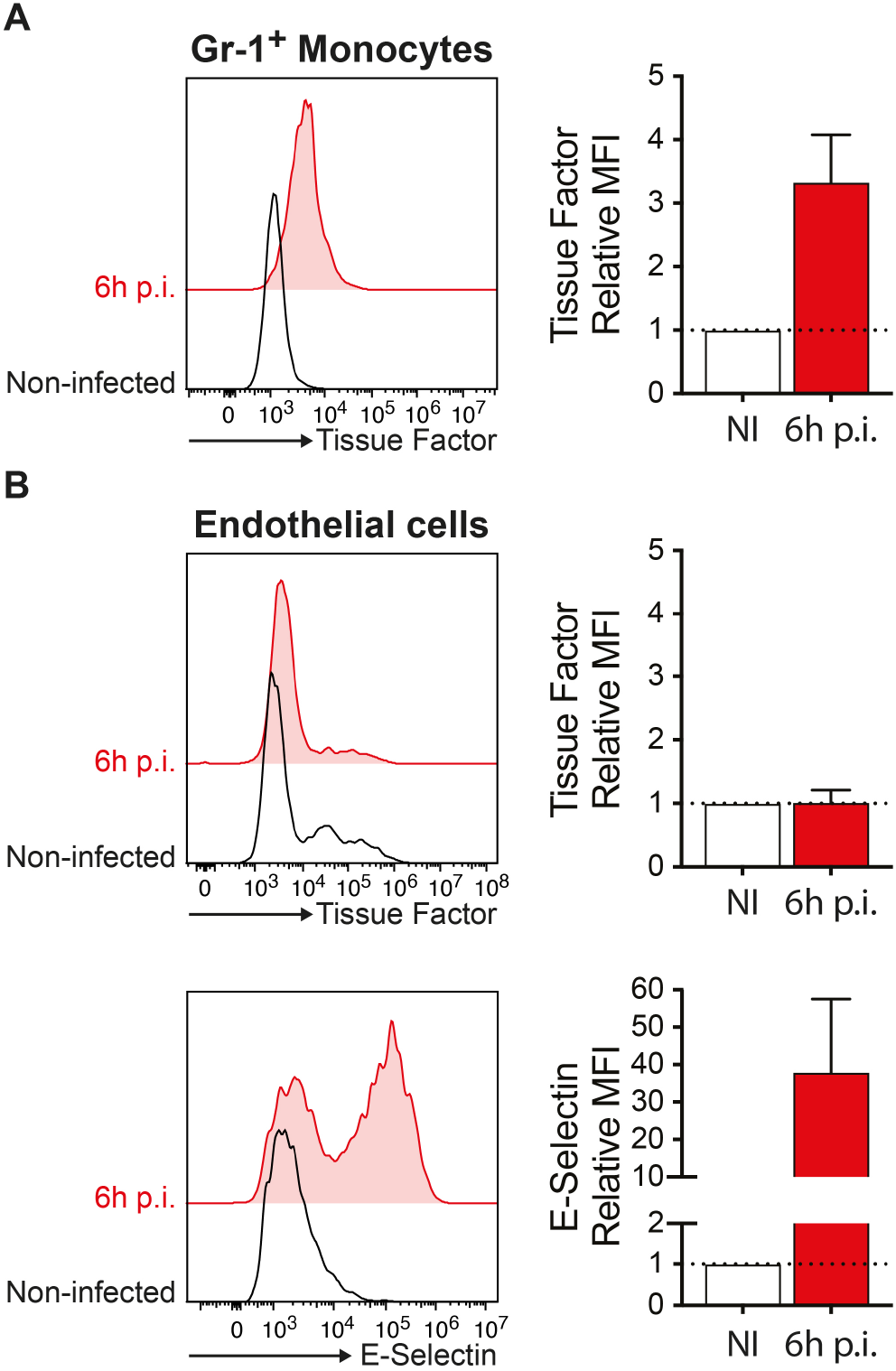
Early upregulation of Tissue Factor at the surface of circulating monocytes upon meningococcal vascular colonization. **A** Representative flow cytometry analysis (left) and quantification (right) of the cell surface expression of murine Tissue Factor (CD142) on blood-circulating monocytes (gated as CD45^+^ CD11b^+^ CD115^+^ Gr-1^+^) in non-infected control mice (black) and mice infected for 6h (red). (n = 3 mice). **B** Representative flow cytometry analysis (left) and quantification (right) of the cell surface expression of human Tissue Factor (CD142, Top) and human E-Selectin (CD62E, bottom) on CD31^+^ human endothelial cells isolated from human skin xenografts in non-infected control mice (black) and mice infected for 6h (red). (n = 3 mice). Data information: In (A-B), data are presented as the mean ± SEM and were normalized with respect to the mean fluorescence intensity (MFI) of the non-infected controls.

## Movie legends

**Movie EV1. Altered blood flow during *Neisseria meningitidis* infection**

High-speed (50 frames/sec) intravital imaging (single focal plane) of 1 μm fluorescent microspheres (white) intravenously perfused allowing the visualization and quantification of the blood flow in human blood vessels (UEA1 lectin, gray) at the indicated time points in infected and non-infected control mice. The extend of vascular colonization by iRFP-expressing *Neisseria meningitidis* (green) is shown as a maximum intensity z-projection. Scale bar: 50 μm.

**Movie EV2. Dynamics of platelets during *Neisseria meningitidis* infection**

Simultaneous dual color intravital imaging (single focal plane) of iRFP-expressing *Neisseria meningitidis* (green) and platelets (anti-GPIbβ, magenta) in a human vessel (white dashed line) 2h post infection. Scale bar: 50 μm.

**Movie EV3. Intravascular fibrin deposits in *Neisseria meningitidis* infected human vessels**

3D volume rendering of a fibrin deposit (anti-Fibrin, red) in a human vessel (UEA-1 lectin, gray) infected with iRFP-expressing *Neisseria meningitidis* (green) for 5 hours. Scale bar: 30 μm.

**Movie EV4. Coagulation inhibition restores the blood flow in *Neisseria meningitides* infected vessels**

High-speed (50 frames/sec) intravital imaging (single focal plane) of 1 μm fluorescent microspheres (white) intravenously perfused allowing the visualization and quantification of the blood flow in human blood vessels (UEA1 lectin, gray) before (t0h00) and after (t4h00) infection upon anticoagulant (Fondaparinux) treatment. The extend of vascular colonization by iRFP-expressing *Neisseria meningitidis* (green) is shown as a maximum intensity z-projection. Scale bar: 50 μm.

## Notes

### Competing Interest Statement

The authors have declared no competing interest.

